# Dynamic 2′-O-Me of rRNA regulates ribosome translation in Arabidopsis nucleolin1 mutants and during seed-to-seedling transition

**DOI:** 10.64898/2026.10.05.756201

**Authors:** Sara Alina Neumann, Jacinthe Azevedo-Favory, Marie-Christine Carpentier, Rémy Merret, Virginie Marchand, Valerie Bourguignon, Pascale Comella, Anne de Bures, Magali Rompais, Laurence Drouard, Christine Carapito, Christine Gaspin, Yuri Motorin, Julio Sáez-Vásquez

## Abstract

Across all kingdoms of life, 2′-*O*-ribose methylation (*2*′*-O-*Me) is the most abundant modification of ribosomal RNAs (rRNAs), which are central to ribosome assembly and function. The 2′-*O*-Me of specific rRNA sites is catalysed by C/D-box small nucleolar ribonucleoprotein (snoRNP) complexes. However, how and when the landscape of plant rRNA 2′-*O*-Me modifications is remodelled remains largely unexplored. To study the regulation of rRNA 2′-*O*-Me and its impact on ribosome translation, we characterised Arabidopsis NUCLEOLIN 1 (NUC1)-depleted plants, which exhibit reduced 2′-*O*-Me at dozens specific rRNA sites, and investigated the impact of 2′-*O*-Me in ribosome translation during plant development. We demonstrated that 2′-*O*-Me is dynamically regulated during seed-to-seedling transition. Our analysis identified differentially methylated rRNA sites, which are specifically located in structurally critical regions of the ribosome. Notably, this hypomethylation is uncoupled from the global expression, assembly, or subcellular localisation of C/D-box snoRNPs. We further observed that polysomes with low 2′-*O*-Me levels not only accumulate but also exhibit differential mRNA loading. These polysomes characterised by reduced 2’-*O*-Me at specific rRNA sites, influence multiples aspects of translation, including ribosomes stalling at the initiation, elongation and termination steps. Together, these findings advance our understanding of the dynamic regulation of rRNA 2′-*O*-methylation and its functional role in ribosome-mediated translation in plants.

## INTRODUCTION

Ribosome synthesis and mRNA translation activity are essential for growth and development in all organisms. Ribosome synthesis requires the assembly of ribosomal RNA (18S, 5.8S, 25S and 5S rRNA) with ribosomal proteins (RP) to form small 40S and large 60S ribosomal particles (reviewed in (1–4)). The keystone of ribosome synthesis is the transcription of 45S ribosomal DNA genes (45S rDNA), which are located in hundreds of copies within nucleolar organiser regions (NORs). 45S rRNA genes from active NORs are transcribed in the nucleolus by RNA polymerase I as a large 45S rRNA precursor (45S pre-rRNA) comprising the rRNAs 18S, 5.8S and 25S separated by external (5’ETS and 3’ETS) and internal (ITS1 and ITS2) transcribed spacers. Simultaneously, specific endo- and exonuclease activities remove the ETS and ITS from 45S pre-rRNAs to produce mature 18S, 5.8S and 25S rRNA. The 5S rRNA is transcribed by RNA polymerase III (5).

In *Arabidopsis thaliana*, the initial endonucleolytic cleavage of the 45S pre-rRNA involves the nucleolin-U3 snoRNP protein complex and occurs at the P site (the equivalent of yeast A0 and mammalian 01(A’)) in the 5’ETS, generating the 35S pre-rRNA (5). The resulting 35S pre-rRNA can then be processed via two alternative pathways. In the major ITS1-first pathway, the 35S is first cleaved at the A3 site (in the ITS1) and then at the P’ and P2 sites (in the 5’ETS), whereas in the minor 5’ETS-first pathway, the 35S intermediate is first cleaved at the P’ and P2 sites (in the 5’ETS), and then at the A2 site (in the ITS1) (6,7). Thus, the major ITS1-first and minor 5’ETS-first processing pathways in Arabidopsis are analogous to the major processing pathways used in mammalian cells and yeast, respectively (reviewed in (4,8,9)).

In addition, throughout transcription of the 45S pre-rRNA, the 18S, 5.8S and 25S rRNA sequences undergo site-specific rRNA modifications (reviewed in (4,10,11)). The most abundant rRNA modifications are ribose 2’-*O-*methyation (2’-*O*-Me) and pseudouridylation (Psi), which are found in most domains of the 25S, 18S and in the 5.8S rRNAs, but not in the 5S rRNA (12–16). The 2′-*O*-Me is carried out by C/D box snoRNP complexes containing the methyltransferase FIB/Nop1, whereas Psi is catalysed by H/ACA box snoRNP complexes containing the pseudouridine synthase DKC1/dyskerin (17–19). Other modifications are base methylations (m^6^A, m^1^A, m^5^C, m*^7^*G) and acetylation (ac^4^C) at specific rRNA sites mainly located in the decoding center (DC) and peptidyl transferase centre (PTC) (11,20–22). Altogether, these rRNA modifications may exert structural, functional or still not fully understood roles.

In animal cells, alterations in ribosome 2’-*O*-Me have been linked to cancer and development. Ribosomes with altered 2’-*O*-Me profiles translate mRNA with reduced fidelity and increase internal ribosome entry site-dependent translation initiation of key cancer genes in cells without active tumour suppressor p53 (11,23–25). Furthermore, differential expression of C/D-box snoRNA has been reported in blood serum and plasma (26), and changes in rRNA ribose methylation have been observed in developing mouse tissues (27).

In *A. thaliana*, hundreds of C/D-box snoRNAs have been identified *in silico*, and the expected methylation sites have been localised on the 18S, 5.8S and 25S rRNA sequences (28–32). More recently two independent studies have mapped a total of 118 Nm sites. Four of these Nm were differentially detected in 9- and 21-day-old Arabidopsis plants. (12,33). Compared to human and yeast, 51 Nm appear to be plant-specific, while a subset of 36 Nm sites are conserved in both yeast and human; five Nm sites are conserved only in yeast and 28 Nm only in human (34).

The current data also suggest a role for 2’-*O-*Me in plant growth and development. Disruption of single C/D-box snoRNA genes in Arabidopsis resulted in root and shoot growth defects in *snoR28* plants (35). Furthermore, snoR28 plants, which exhibit reduced 2’-*O*-Me at a single Nm site (25S-Gm2396) show impaired pre-rRNA processing, altered polysome profiles, and defects in root and shoot growth (Cao et al., 2022). Notably, SnoR28.1 and NUC1 interact physically and genetically, however the Nm site targeted by SnoR28.1 is not hypomethylated in *nuc1* (12). Similarly, disruption of both the snoRNP assembly factor gene Nufip/Nrsa and the nucleolar protein NUC1, resulted in reduced rRNA 2’-*O*-Me (12,36–39). Despite these observations, the underlying regulatory mechanisms controlling 2’-*O*-Me and their impact on ribosome activity remains only partially understood. To address this question, we further characterise C/D-box snoRNP expression and assembly in *nuc1* plants and analyse rRNA 2’-*O*-Me during Arabidopsis seed-to-seedling transition.

## RESULTS

### 2’-*O*-Me of specific Nm sites and Expression of C/D-box snoRNPs are not co-regulated

We previously reported reduced 2’-*O*-methylation in Arabidopsis nucleolin 1 (NUC1) knock-out plants (12). In contrast, the NUC2 protein, which accumulates in *nuc1* plants (37,40) and during early germination (41), does not appear to be involved in 2’-*O*-Me of rRNA. To determine whether the reduced rRNA 2’-*O*-Me levels observed in *nuc1-2* plants depend on the differential expression of the C/D-box snoRNP complex(es), we analysed the accumulation of FIB and NOP56 proteins, as well as C/D-box snoRNAs (**Fig. 1**).

**Figure 1:**
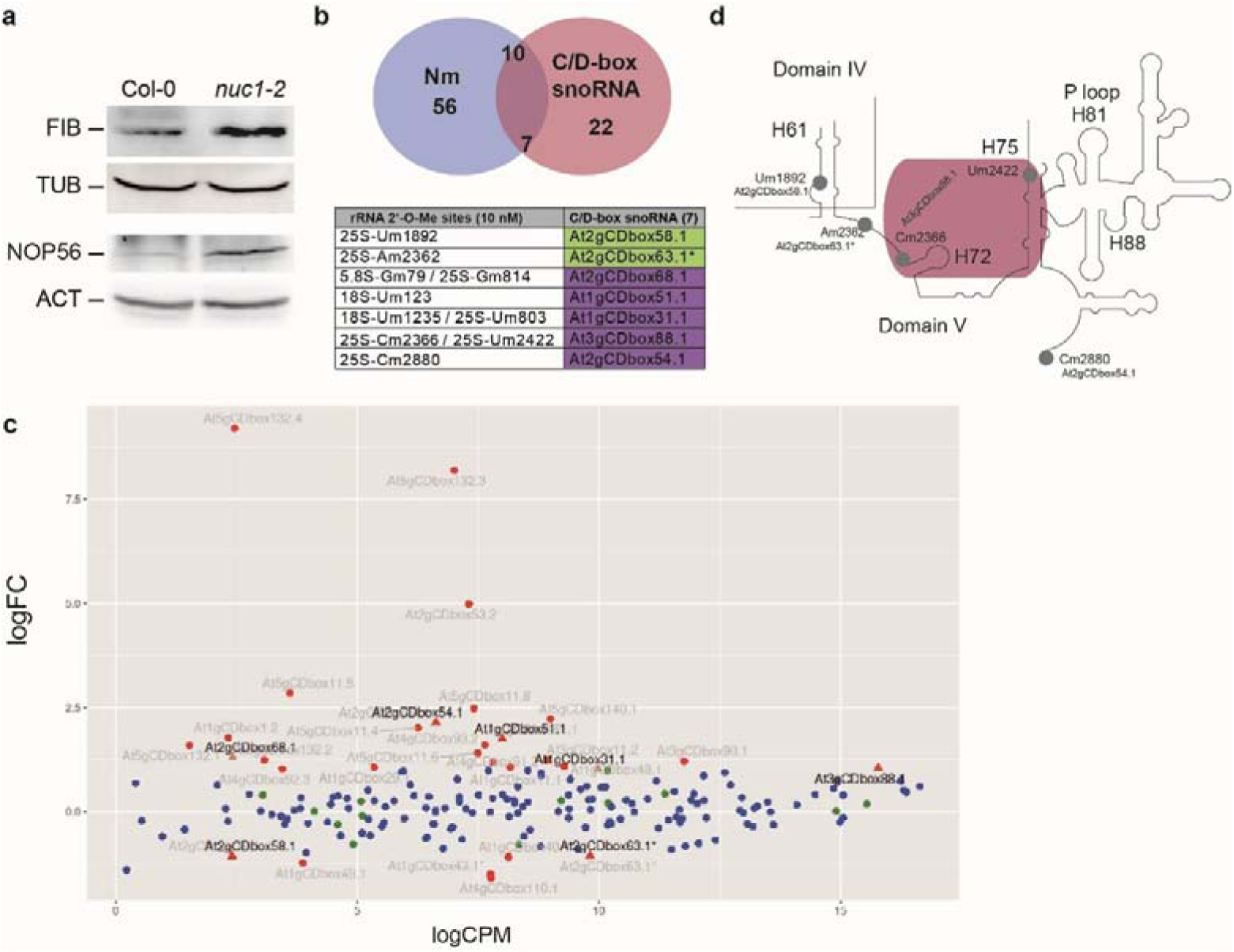
rRNA 2’-*O*-Me and C/D-box snoRNP expression. **a**, Immunodetection of FIB (FIB1, 32.83 kDa and FIB2, 33.65 kDa) and NOP56 (At3g12860, 55.4 kDa and At1g56110, 57.9 kDa) isoforms in 21-day-old soil-grown Col-0 and *nuc1-2* plants using α-38F3 and α-NOP56. α-TUBULIN (TUB) and α-ACTINE (ACT) were used to verify amounts of proteins in each sample. **b**, RNA-seq analysis of small RNA fraction (< 200 bases) and analysis of C/D-box snoRNAs variations in biological triplicates of 18-day-old (soil-grown) Col-0 and *nuc1-2* plants. The Venn diagram shows 66 Nm (light blue; *p*.adj ≤ 0.05; Cohen’s *d* ≥ 0.8) and 29 C/D-box snoRNAs with significant changes (pink; *p* ≤ 0.01) in *nuc1-2* compared to Col-0. Ten hypomethylated Nm are associated with changes in seven C/D-box snoRNA accumulation. The table shows the 10 hypomethylated Nm, the two down-regulated C/D-box snoRNA showing positive correlation (green) and the five up-regulated C/D-box snoRNA with negative correlation (purple). The asterisk in At2gCDbox63.1 indicates a snoRNA that is dicistronic with a tRNA. **c,** Minus average (MA) plot showing the logarithm of the fold-change (logFC) and the average log count (logCPM) of C/D-box snoRNA in *nuc1-2* compared to Col-0. Coloured dots show non-significant (blue) and significant (red) differential expression (DE) of C/D-box snoRNAs. C/D-box snoRNAs with logFC ≥ 1.5 or logFC ≤ −1.5 were labelled and the seven significantly differentially expressed C/D-box snoRNAs correlating with 10 hypomethylated Nm were indicated with a triangle instead of a dot. **d,** Scheme shows the location of 2’-*O*-Me target sites of four C/D-box snoRNAs differentially expressed. Grey circles indicate rRNA Nm, whose corresponding C/D-box snoRNAs show significant differences in their expression between Col-0 and *nuc1-2*. Purple box mark two rRNA sites targeted by At3gCDbox88.1. The asterisk in At2gCDbox63.1 indicates a snoRNA that is dicistronic with a tRNA.

Western blot analysis detected slightly elevated FIB and NOP56 signals in *nuc1-2* compared to Col-0 (**Fig. 1a**). FIB1 and FIB2, as well as the two NOP56 orthologues, have similar molecular weights. Consequently, neither the two FIB nor the two NOP56 orthologous proteins can be distinguished from each other by Western blot. Nevertheless, transcriptomic analysis of mRNAs from Col-0 and *nuc1-2* plants revealed a slight accumulation of transcripts encoding all snoRNP core proteins.

Then, we performed RNAseq analysis of small RNA fractions from Col-0 and *nuc1-2* plants (**Figs. 1b-c**). The Principal Component Analysis (PCA) shows variance between Col-0 and *nuc1-2*, with clear clustering of biological triplicates. Approximately 20M reads, longer than 40 nt, were obtained by sample after cleaning reads. FlagStats metrics of mapping show that 22.82% to 44.96% of these reads aligned on C/D-box snoRNAs for all samples. Of these, 29 C/D-box snoRNAs (over the 143 targeting the 117 rRNA 2’-*O*-methylation sites (Azevedo-Favory et al., 2021)) showed differential accumulation in *nuc1-2* compared to Col-0 plants. However, only seven C/D-box snoRNA with significantly altered expression levels targeted 10 Nm sites showing noticeable change in rRNA 2’-*O*-Me levels in *nuc1-2* compared to Col-0 (**Fig. 1b**). Two of these C/D-box snoRNAs were significantly downregulated and their corresponding Nm sites showed significant hypomethylation compared to Col-0 (At2gCDbox58.1/25S-Um1892 and At2gCDbox63.1*/25S-Am2362). In the other five cases (At2gCDbox68.1, At1gCDbox51.1, At1gCDbox31.1, At3gCDbox88.1, At2gCDbox54.1) an upregulation of C/D-box snoRNA levels was observed, while the corresponding Nm was significantly hypomethylated. The two Nm sites showing positive correlation between 2’-*O*-Me level and C/D-box snoRNAs expression are located in domains IV and V of the 25S rRNA sequence while two other Nm sites are targeted by a single C/D-box snoRNA (**Figs. 1d**).

We conclude that global changes in 2’-*O*-Me levels in *nuc1-2* are unlikely to be regulated by alterations in C/D-box snoRNP expression.

### C/D-box snoRNPs are found in disorganised nucleoli

Catalytically active C/D-box snoRNPs are targeted to nucleoli where they function in 2’-*O*-Me of rRNAs (17). In *nuc1-2* plants, the functional organisation of the nucleolus is disrupted (37) and induces changes in the distribution of nucleolar and non-nucleolar proteins (42). Therefore, we investigated subcellular localisation of C/D-box snoRNPs in Col-0 and *nuc1-2* seedlings.

We used a FISH (Fluorescence *in situ* hybridisation) probe mix for seven C/D-box snoRNAs that drive 2’-*O*-Me and the FISH probe for U3 snoRNA that is involved in rRNA cleavage. In both Col-0 and *nuc1-2,* the C/D-box snoRNAs (including U3) involved in 2’-*O*-Me or in rRNA cleavage are located in the nucleolus as judged by the overlay of a FISH signal and DAPI staining. RNase treatment of Col-0 and *nuc1-2* samples makes the FISH signals disappear, indicating specific detection of C/D-box snoRNAs in the nucleolus (**Figs. 2a**).

**Figure 2:**
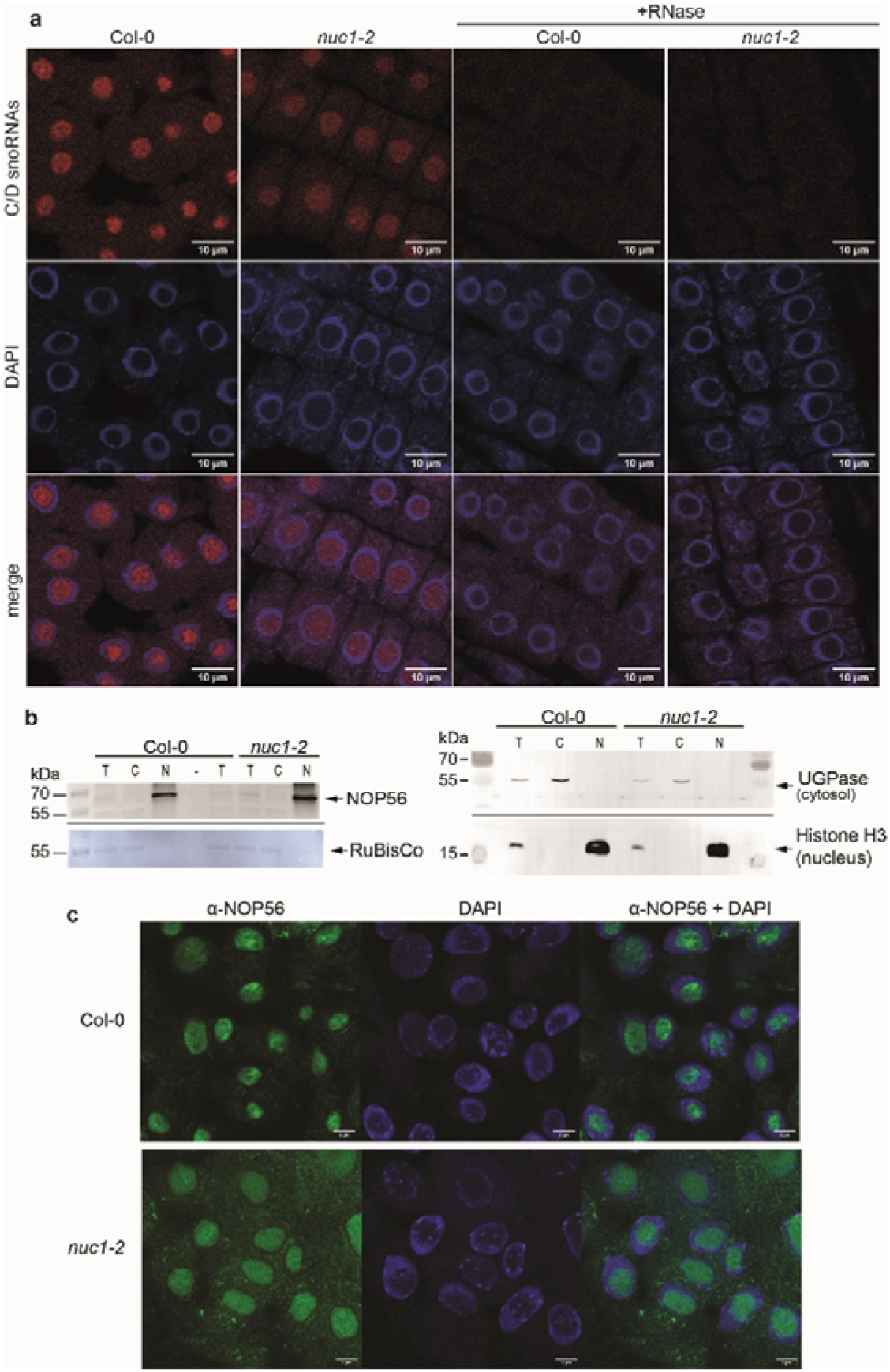
Localisation of C/D-box snoRNPs in functionally disorganised nucleoli. **a,** FISH experiment on root cells from 7-day-old (*in vitro*-grown) Col-0 and *nuc1-2* seedlings using 3’-cyanine labelled probes for C/D-box snoRNAs (At5gCDbox137.1, At1gCDbox25.1/HID2, At1gCDbox33.1, At5gCDbox141.1, At4gCDbox117.1 and At1gCDbox18.1). The FISH probes reveal snoRNAs (red) in the nucleoli of Col-0 and *nuc1-*2. FISH experiments before (-RNase) and after (+RNase) RNase treatment are shown. Counterstaining of FISH with DAPI (blue) is shown (merge). Scale bar set at 10 µm. **b,** Western blot of NOP56 in total (T), cytosolic (C) and nuclear (N) fractions from 20-day-old (*in vitro* grown) Col-0 and *nuc1-2* seedlings. The Coomassie blue-stained membrane after hybridisation with α-NOP56 is shown (left) together with immuno-detection of UGPase and histone H3, which are a cytosolic and a nuclear protein respectively; and used to control T, C and N fractions (right panel). **c,** Bottom panels, immunolocalisation of NOP56 (green signal) in the nucleolus of Col-0 and *nuc1-2* plants. DAPI staining (blue) was used to visualise the nucleoplasm and distinguish the nucleolus. Scale bar set to 5 µm.

We then determined the subcellular localisation of core protein NOP56. Western blot analysis revealed a strong NOP56 signal in the nuclear fractions of both Col-0 and *nuc1-2* plants, with a weaker signal in the total but none in the cytosolic fraction (**Fig. 2b**). Immunolocalisation further confirmed NOP56 primarily in the nucleolus of both Col-0 and *nuc1-2* plants (**Fig. 2c**). Similarly, fibrillarin is detected in the nuclear fraction and in the disorganised nucleolus of *nuc1-2* as previously described (40).

Overall, these results suggest that rRNA hypomethylation in *nuc1-2* plants is unlikely to be due to a failure of nucleolar localisation of C/D-box snoRNA or core protein factors.

### Fibrillarin and NOP56 co-purify with C/D-box snoRNA in *nuc1-2* plants

One possible explanation for the reduced rRNA 2’-*O*-Me in *nuc1-2* plants is the failure to assemble a catalytically active C/D-box snoRNP complex. To investigate this, we used Col-0 and *nuc1-2* plants expressing FIB2-YFP, which localises to the nucleolus (42).

Expression of FIB2-YFP does not alter Col-0 or *nuc1-2* phenotypes (**Fig. 3a**) and accumulates similarly FIB2-YFP in both genetic backgrounds. However, endogenous fibrillarin levels remain slightly elevated in *nuc1-2*+FIB2-YFP versus Col-0+FIB2-YFP (**Fig. 3b**), consistent with untransformed lines. LC-MS/MS analysis revealed a significant accumulation of NOP58 (At5g27120, LogFC 1.9, *p*-value 0.0076) and NOP56 (At1g56110.1, LogFC 0.79, *p-*value 0.0042) peptides in *nuc1-2*. No significant changes were detected for FIB2 peptides (LogFC 0.65, *p-*value 0.26), probably due to the FIB2-YFP overexpression. Peptides for the NOP58 orthologue At3g05060 (LogFC 1.6, p-value 0.086) and the L7Ae orthologues AT2G47610 (LogFC −0.55, p-value 0.22) and At3g62870 (LogFC −0.89, p-value 0.045) were detected at similar levels in both Col-0 and *nuc1-2* backgrounds in both backgrounds. This analysis also revealed reduced accumulation of 60S ribosomal proteins RPL34e/eL34x and RPP3A/P3z, but increased accumulation of LA1, the Lhp1p orthologue that binds and stabilises C/D-box U3 snoRNA precursors (43).

**Figure 3:**
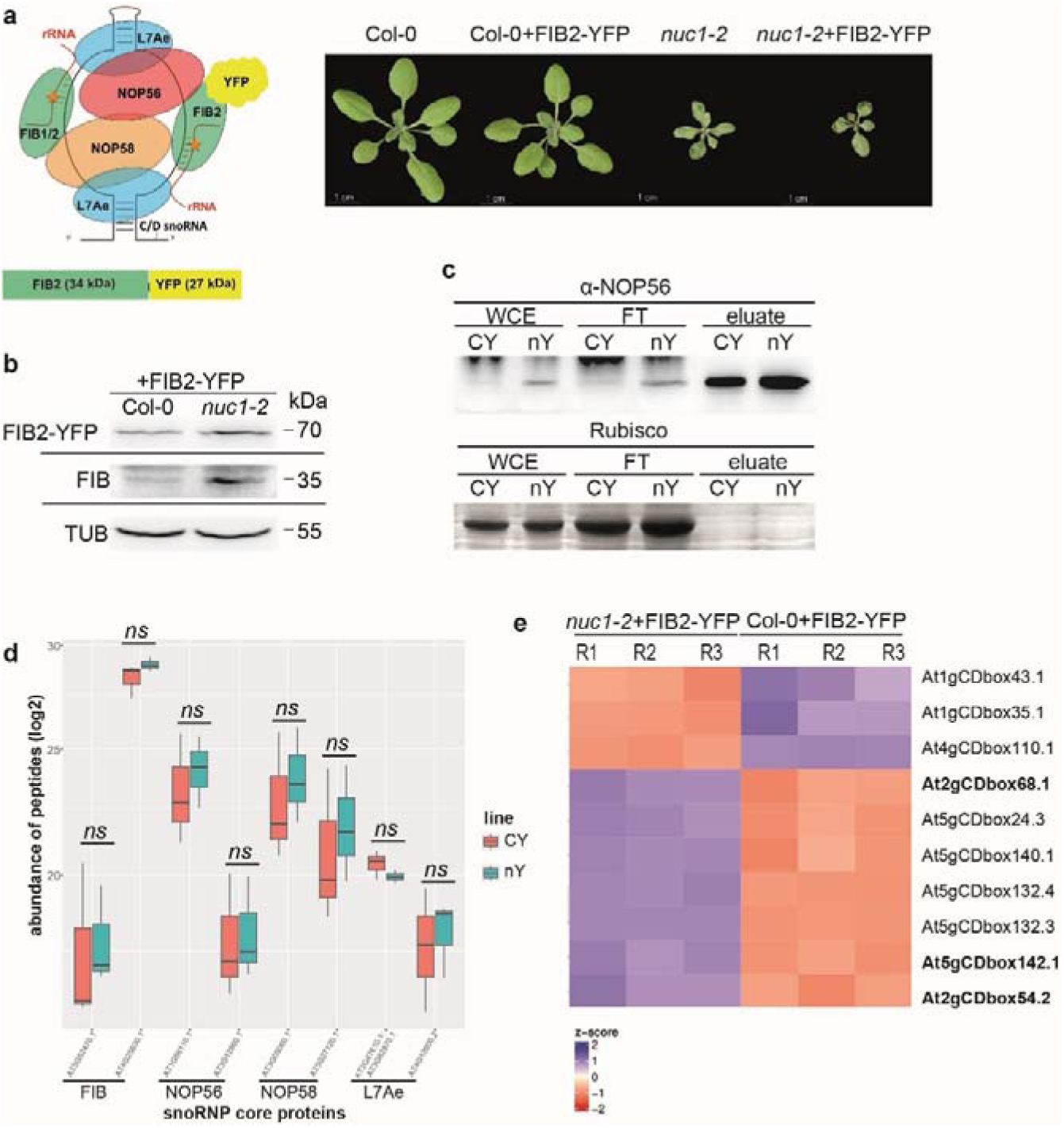
Expression and assembly of C/D snoRNP core proteins. **a**, Left, schematic representation of the C/D-box snoRNP complex and the FIB2-YFP chimeric protein of expected size of 61 kDa. Right, plant phenotypes for 21-day-old (soil-grown) Col-0 and *nuc1-*2 plants expressing or not FIB2-YFP. **b**, Western blot using α-38F3 to detect FIB2-YFP and FIB1/FIB2 (∼35 kDa) in 21-day-old (soil-grown) Col-0+FIB2-YFP and *nuc1-2*+FIB2-YFP plants. Since FIB2-YFP expression is much stronger than endogenous FIB1/FIB2 protein expression, FIB2-YFP and endogenous FIB1/2 signals are shown separately with different contrasts. TUBULIN was used as a loading control to insure the similar amount of protein in each sample. **c**, Immunoprecipitation (IP) of C/D-box snoRNP proteins using α-GFP and Col-0+FIB2-YFP (CY) and *nuc1-2*+FIB2-YFP (nY) whole cell extracts (WCE). Top, Western blot of WCE, flow through (FT) and eluate fractions with α-NOP56. Bottom: Coomassie blue staining after immunodetection showing the Rubisco band to verify similar protein loading. **d**, Bar chart shows accumulation of orthologues FIB1/FIB2, NOP56, NOP58 and L7Ae detected by LC-MS/MS in the IP eluate fractions from *nuc1-2*+FIB2-YFP (nY) compared to Col-0+FIB2-YFP (CY) (3 replicates). Welch’s t-test does not detect significant difference in the accumulation of FIB1/FIB2, NOP56, NOP58 and L7Ae peptides. ns not significant value ≥0.05. **e,** Heat map representation of snoRNA enrichment in the IP eluated fractions from three Col-0+FIB2-YFP and *nuc1-2*+FIB2-YFP biological replicates. The rRNA sites are clustered according to hclust/ward.D2 method. The colour key, histogram and values are shown. Bold, indicates C/D-box snoRNAs associated with reduced 2’-O-Me in *nuc1-2* plants.

Then, we performed immunoprecipitation (IP) of FIB2-YFP in *nuc1-2* and Col-0 plant lines to investigate failure of core proteins to interact with each other and/or with C/D-box snoRNAs (**Fig. 3c**). Strong NOP56 signals were detected in eluates from both *nuc1-2*+FIB2-YFP and Col-0+FIB2-YFP, indicating FIB2-YFP and NOP56 interaction in both genotypes (**Fig. 3c**). LC-MS/MS analysis identified 873 proteins in the eluates, with similar peptides abundance for FIB1/2, NOP56, NOP58 and L7Ae orthologues in both *nuc1-2*+FIB2-YFP and Col-0+FIB2-YFP (**Fig. 3d**).

We also performed RNAseq analysis on eluates from *nuc1-2*+FIB2-YFP and Col-0+FIB2-YFP, detecting 196 snoRNA in both fractions, with a good reproducibility between genotypes. Seven C/D-box snoRNAs were differentially accumulated in *nuc1-2+FIB2-YFP* eluates. For three of these, the 2’- *O*-Me levels at their corresponding Nm sites was reduced, while the 2’-*O*-Me level of the Nm target sites for the other snoRNAs remained unaffected. The three C/D-box snoRNAs reduced in *nuc1-2*+FIB2-YFP eluates do not have Nm target sites in the rRNA (**Fig. 3e**).

Taken together, these results indicate that the synthesis and interactions of the C/D-box snoRNP core proteins and snoRNAs are not compromised in *nuc1-2* plants.

### Hypomethylation of rRNA / ribosomes provokes polysomes accumulation in *nuc1-2* plants

To investigate how rRNA hypomethylation at specific Nm sites in *nuc1-2* plants affects translational activity, we first performed polysome profiling in Col-0 and *nuc1-2* plants. In *nuc1-2*, a significant accumulation of polysomes and a slight reduction of monosomes (80S) was observed compared to Col-0 (**Fig. 4a**, left panel). Quantification of the areas under the curves for the polysome profiles showed that both the polysome/80S monosome ratio and the percentage of polysomes relative to the total profile (40S, 60S, 80S and polysomes) were higher in *nuc1-2* (**Fig. 4a**, right panel). The accumulation of polysomes and 2’-*O*-Me profiles in *nuc1-2* plants transformed with a 35S::EGFP-NUC1 or 35S::NUC1-EGFP construct was similar to that observed in Col-0 plants, indicating that the EGFP tagged NUC1 proteins are able to rescue disruption of the endogenous NUC1 gene.

**Figure 4:**
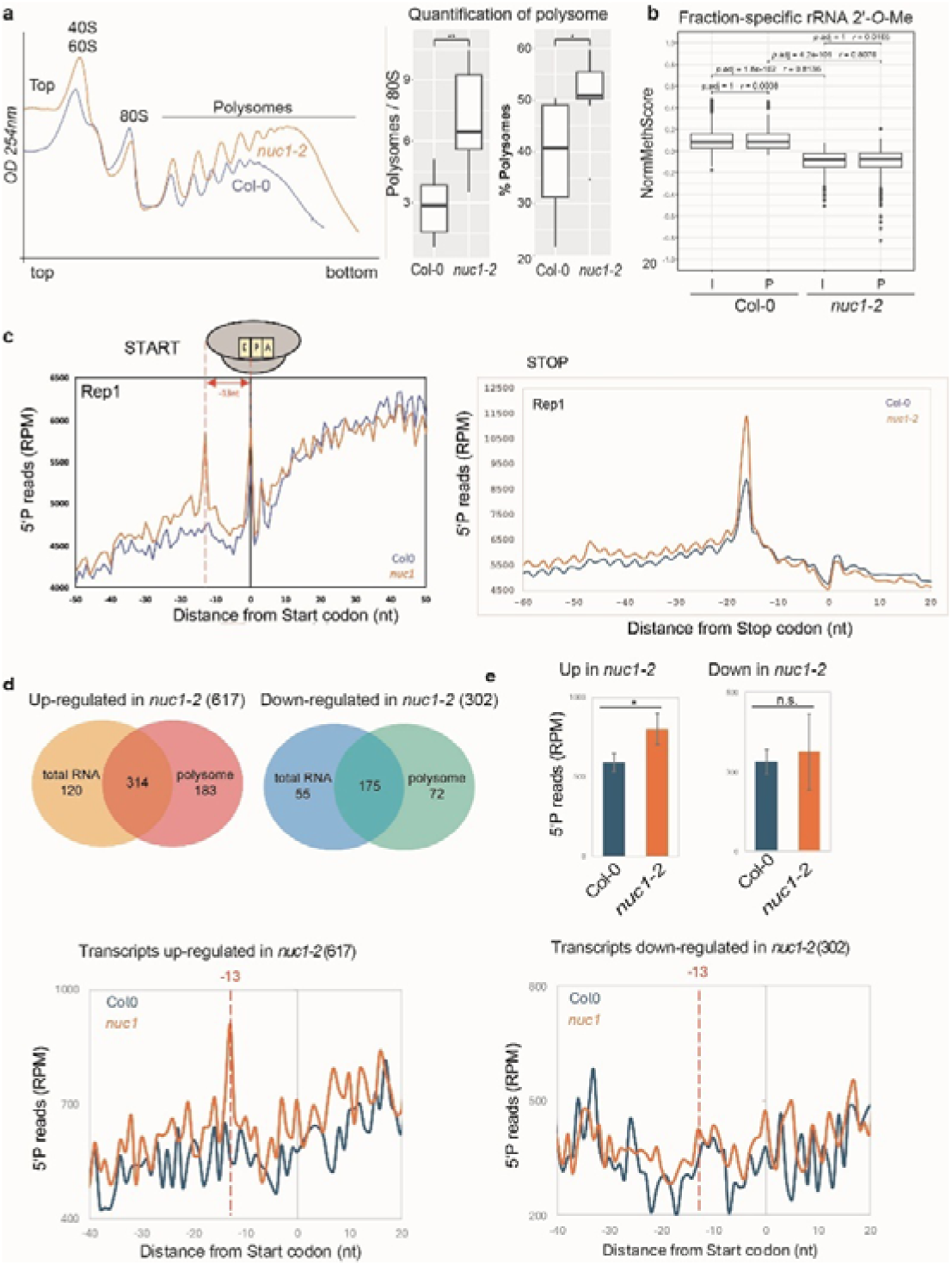
Depletion of NUC1 impacts polysome profile and translation. **a**, Polysome profiles of 18-day-old (soil-grown) *nuc1-2* (red) and Col-0 (blue) plants and quantification in six biological replicates of the polysome/80S area ratio and the percentage (%) of polysomes out of the total area of 40S, 60S, 80S, polysomes. **b**, Box plots of the average NormMethScore across all 117 rRNA 2’-*O*-Me sites in biological triplicates of input (I) and polysome (P) fraction from Col-0 and *nuc1-2*. Statistical significance of rRNA 2’-*O*-Me patterns was determined using Wilcoxon test and the resulting *p*-values were adjusted using the Holm-Bonferroni method. Additionally, Wilcoxon effect size was used to compute non-parametric effect size *r*. Comparisons with *p*.adj <0.05 and an effect size *r* ≥0.3 were considered significantly different. **c**, Metagene analysis of 5’P sequencing (5’Pseq) approach as previously described in (47,48) displaying the abundance of 5‘P reads in rpm relative to the start, stop and CCG, TGC, TGT, ATA, TTG and TGG codons in *nuc1-2* (red) compared with Col-0 (blue) plants grown for 18 days on soil. Representation of the ribosome during elongation shows the 13 nt distance between the 5‘P peaks and the P site of the ribosome. **d**, Venn diagrams show up- and down-regulated RNA transcripts in total and polysome RNA fractions from Col-0 and *nuc1-2* plants. **e,** Metagene analysis around the start codon of transcripts significantly upregulated (A) or downregulated (B) in *nuc1* compared to Col0 in total RNA-seq and/or Polysome RNA-seq. Histograms represent read accumulation at −13 nt upstream of the start codon in each genotype. N = three biological replicates. *= *p* < 0.05 and ns not significant value *p* ≥0.05

Next, to test whether these differentially accumulated polysomes showed any bias in the rRNA 2’-*O*-Me pattern, we extracted RNA from the total protein/RNA extracts and the polysome fraction of Col-0 and *nuc1-2* and performed RiboMethSeq analysis. We detected 117 Nm: 38 in the 18S rRNA, two in the 5.8S rRNA and 77 in the 25S rRNA sequences (**Fig. 4b**). On a global level, the results show statistically significantly (*p*.adj ≤ 4.2e-101) decreased 2’-*O*-Me levels in *nuc1-2* both input and polysome fraction compared to the respective fraction in Col-0 (*r* > 0.8) (**Fig. 5b**). Notably, 92 significantly (*p*.adj < 0.05 & *d* ≥ 0.8) hypomethylated Nm sites in the *nuc1-2* input were also significantly hypomethylated in the *nuc1-*2 polysome fractions. A further four and eight sites, respectively, were significantly hypomethylated in *nuc1-2* polysomes but not in input fractions, or conversely hypomethylated in *nuc1-2* input but not in *nuc1-2* polysome fractions.

**Figure 5:**
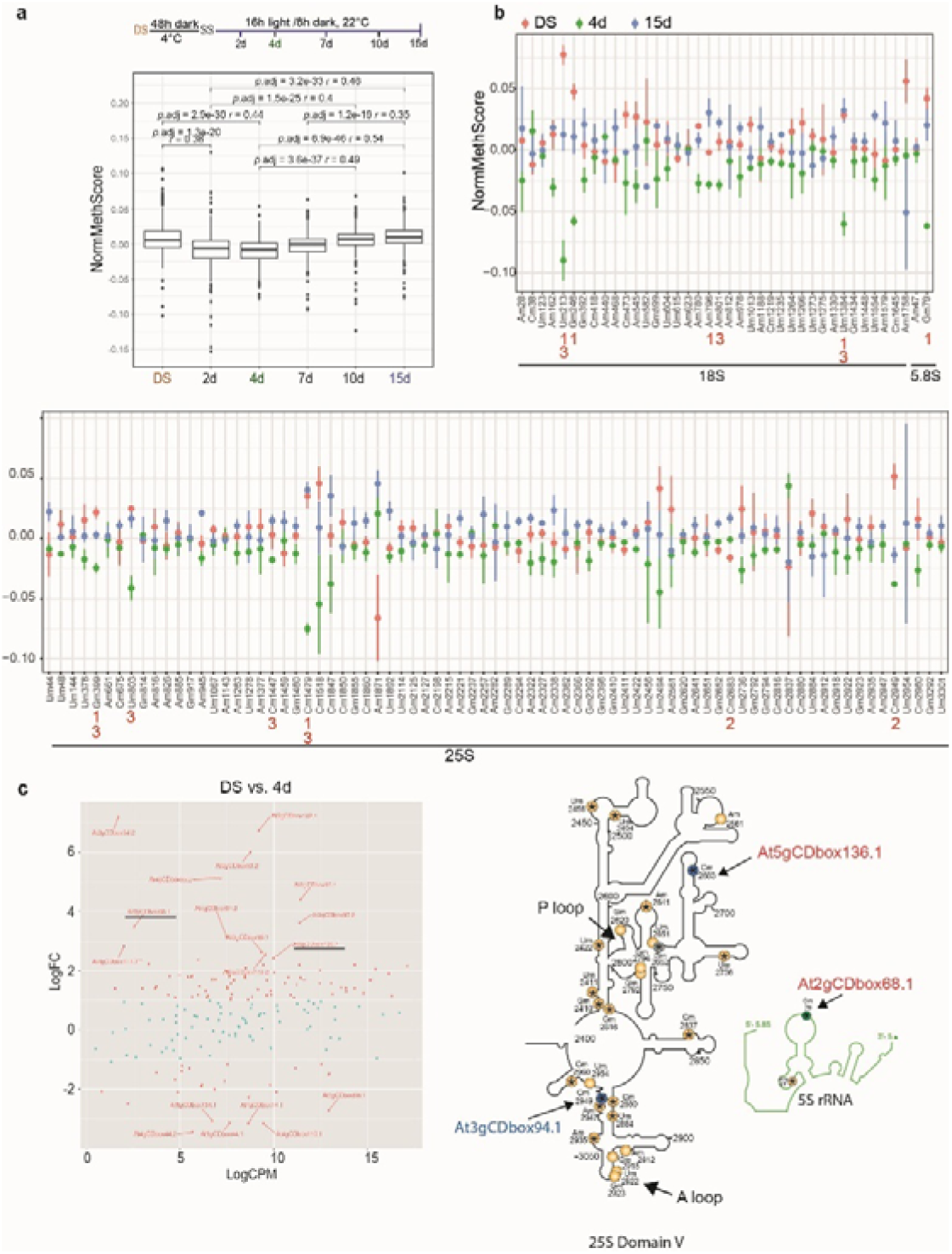
rRNA 2’-*O*-Me patterns during early development in Arabidopsis. **a**, Experimental design to obtain 2’-*O*-Me, polysome and RNA-seq data. Box plot of the average normalised MethScore values across 117 rRNA 2’-*O*-Me sites in three biological triplicates of Col-0, DS, 2d, 4d, 7d, 10d and 15d. Statistical significance was determined by Wilcoxon test and the resulting p-values were adjusted using the Holm-Bonferroni method (*p*.adj ≤ 1.2e-19). Additionally, Wilcoxon effect size was used to compute non-parametric effect size r (*r* ≥ 0.3). Only significant comparisons are indicated (p.adj < 0.05 & r ≥ 0.3) **b,** Dot plots show Nm Normalised MethScores (NormMethScore) with error bars of all 117 Nm sites in the 18S/5.8S (top panel) and 25S (bottom panel) rRNA in biological triplicates of Col-0 DS, 4- and 15-day-old seedlings. Statistical significance between DS, 4- and 15-day-old seedlings was verified using Welch’s t-test adjusted using the Benjamini-Hochberg method. Additionally, standardised mean difference was calculated using the Welch’s compatible Cohen’s *d* formula. Comparisons with *p*.adj < 0.05 and *d* ≥ 0.8 were indicated: DS vs. 4d = 1, DS vs. 15d = 2 and 4d vs. 15d = 3. **c,** MA plots show C/D-box snoRNAs with significant changes in accumulation (logFC ≥ 1.5 or logFC ≤ −1.5) between dry seeds (DS) versus 4-day-seedlings. **d**, Location of three Nm sites, along with their corresponding C/D-box snoRNAs, which are differentially expressed in DS vs. 4-day-old seedlings (green circle in 5S) or in DS vs 15-day-old seedlings (bleu circles in 25S). Yellow circles indicate non-significant (ns) Nm changes, and grey circles Nm sites without snoRNA drivers. Asterisks (*) within the circles denote NUC1 dependency.

We then assessed whether the polysome profile in *nuc1-2* was caused by the loss of interaction between NUC1 and translating ribosomes/polysomes. Western blot analysis showed that an EGFP-NUC1 protein was mainly detected in the free RNA/upper fraction, with weak signals in the 40S/60S/80S and trailing signals in the polysome fractions. Consisting with this, immunoprecipitation of ribosomes using tagged HF-RPL18 plants (44) detect co-elution of ribosomal subunits RPL13 and RPS6 but no NUC1.

We next examined if 2’-*O*-Me defects and polysome accumulation are functionally linked in other ribosome biogenesis factors (RBFs) plant mutants. First, we analysed the *fibrillarin* 1 mutants, which also exhibit reduced 2’-*O*-Me (33). Unlike *nuc1* plants, the *fib1* mutants display no apparent growth or developmental abnormalities, and their polysome profiles remain comparable to wild-type (Col-0) plants. This discrepancy may stem from the higher number of hypomethylated Nm sites and generally lower methylation levels observed in *nuc1-2*. We also examined the 2’-*O*-Me profile of *apum23* mutants, which share *nuc1-2*’s growth, developmental, and pre-rRNA processing defects but lack polysome profile alterations (45,46). No significant changes in 2’-*O*-Me were detected in these.

Collectively, these findings suggest that polysome accumulation in *nuc1-2* is likely a consequence of ribosomal hypomethylation, rather than a loss of NUC1-polysome interaction. Additionally, impaired pre-rRNA cleavage does not appear to affect 2’-*O*-Me.

### Hypomethylated polysomes provokes impaired translation

The observed accumulation of polysomes in *nuc1-2* suggests a potential defect in ribosome dynamics. Therefore, we performed a 5’Pseq approach that provides *in vivo* information of ribosome footprints (47,48). This approach was already used to reveal ribosome dynamics in several contexts (48–52). In fact, this approach captures co-translational decay events (48). The last translating ribosome is followed by the exoribonuclease codon per codon (XRN1 in yeast and XRN4 in Arabidopsis). The first nucleotide of each 5’P read corresponds to the 5’P boundary of the ribosome. If the ribosome is slowing down at a specific location, a clear 5’P accumulation upstream of this location is expected (53). In Arabidopsis, a clear 5’P accumulation 16 nt before the stop codons was observed (54–56). This position corresponds exactly to the distance between the 5’P boundary of the ribosome and the stop codon in the A-site. If a defect occurs in the P-site, a 5’P accumulation is expected 13 nt before (53).

PCA on 5’Pseq data revealed good reproducibility between Col-0 and *nuc1-2* genotypes. In addition, a clear 3-nucleotide periodicity is observed in each genotype revealing that ribosome footprints were correctly captured by 5’Pseq. To obtain an overview of ribosome dynamics in *nuc1-2*, a metagene analysis around start and stop codons was performed. For the start position, a clear 5’P reads accumulation is present 13 nt before the start codon in *nuc1-2* compared to Col-0 (**Fig. 4c left panel**). This peak is observed independently in each biological replicate and corresponds exactly to the position of a ribosome stalled at the initiation step at the P site. A second peak of 5’P reads at 0 (ATG) position shows similar accumulation in both *nuc1-2* and Col-0 plants. This peak was previously observed in Arabidopsis studies and corresponds probably to specific ribosomes dynamics during the first elongation steps (55). In addition to stalling at the initiation step, additional pauses were observed 16 nucleotides before the stop and several other codons (**Fig. 4c right panel**), corresponding to a ribosome stalled at the A-site of these codons. We then examined whether the arrest or inhibition of translational elongation from the ATG start codon or other codons was associated with changes in tRNA expression. Mim-tRNAseq and tRNA expression analysis revealed no significant differences in tRNA expression in *nuc1*-*2*.

To characterise RNA transcripts that accumulate in *nuc1-2* and display a peak at 13 nt upstream of the start codon, we performed RNAseq analysis of total- and polysome-RNA fractions (**Fig. 4d**). PCA on RNAseq data revealed good reproducibility between total and polysome from Col-0 and *nuc1-2* samples. This analysis revealed 664 differentially expressed (DE) genes (Log2 FC −1/+ 1, p value ≤ 0.05) genes (434 up-regulated and 230 down-regulated) and 745 RNA sequences with differential polysome association (498 with higher loading and 247 with lower loading) in *nuc1-2* compared to Col-0. Among these, 184 transcripts with higher polysome loading and 72 with lower loading were not differentially expressed in *nuc1-2* (**Fig. 4d**). Transcripts preferentially accumulating in *nuc1-2* encode proteins involved in RNA metabolism, gene-specific transcriptional regulation, and translation.

Then, to specifically assess the direct impact of hypomethylated and stalled ribosomes on global mRNA translation, we performed a metagene analysis around the start codon on transcripts significantly upregulated or downregulated in *nuc1-2* compared to Col-0 in total RNAseq and/or polysome RNAseq (N = 617 and 301, respectively). The −13 nt peak observed across the whole transcriptome was detected only in the upregulated transcript population, suggesting that transcripts upregulated in *nuc1-2* display defects in translation initiation. This effect appears to be absent in the downregulated population (**Fig. 4e**). Altogether, these results suggest that transcripts upregulated in *nuc1-2*, especially those in polysomal fractions, are likely to accumulate because of translation stalling at multiple stages, particularly during initiation.

Overall, the results show ribosomes stalling at the initiation, elongation and termination steps in *nuc1-2*. Furthermore, the data suggest that the impact of translation in *nuc1-2* is more likely to be related to functional structural changes in ribosomes than to the expression or loading of specific RNA transcripts.

### 2’-*O*-Me is dynamically regulated during seed-to-seedling transition

We previously reported that *nuc1-2* plants exhibit severe developmental defects at the seedling stage, indicating that rRNA methylation plays an essential role during the transition from seed to established seedling (37,40). Consequently, we investigated the 2′-*O*-Me landscape and polysome profiles throughout this critical developmental transition by performing RiboMethSeq and polysome profiling on dry seeds (DS), 2-, 4-, 7-, 10-, and 15-day-old Arabidopsis seedlings following transfer to growth conditions (**Fig. 5**).

RiboMethSeq allowed quantification of a total of 117 Nm residues: 38 in the 18S rRNA, two in the 5.8S rRNA and 77 in the 25S rRNA sequences. The PCA analysis shows that replicates for DS, 2-, 4-, 7-, 10- and 15-day-old seedlings cluster together, respectively, with some variability between conditions/replicates. The NormMethScores and RawMethScores and average of global rRNA 2’-*O*-Me levels of the NormMethScores for all detected rRNA 2’-*O*-Methylated sites (Nm) for each replicate/condition were calculated and compared to each other.

On a global level, the results indicate a moderate but statistically significant (*p*.adj ≤ 1.2e-1 and *r* ≥ 0.3) gradual decrease in 2’-*O*-Me levels from DS to 4-days-old seedlings followed by a continuous increase from 7- to 10-days-old seedlings (**Fig. 5a**). Global 2’-O-Me levels did not differ significantly between 4 and 7 as well as 10- and 15-day-old seedlings (*p.*adj ≤ 2.9e-30 and *r* > 0.44 for all three comparisons), we therefore selected DS, 4- and 15-days-old seedlings samples for site-specific statistical analysis The 2’-*O*-Me levels of 12 specific rRNA sites change significantly (*p.*adj ≤ 0.05 and *d* ≥ 0.8) when comparing Nm in DS, 4-day-old and/or 15-day-old samples: seven sites in DS vs 4-day-old, two sites in DS vs 15-day-old and seven sites in 4-day-old vs 15-day-old (**Fig. 5b**). The 2’-*O*-Me levels of these 12 Nm sites were also affected in the *nuc1-2* plants (12). The 2’-*O*-Me rRNA sites with significant developmental changes are located in the 3’, 5’ or central domain of the 18S rRNA and in the domains I, II, III and V of the 25S rRNA and in the 5.8S rRNA. Also, three Nm (Um213, Gm246 & Am796) in the 18S rRNA and four Nm (Gm399, Um803, Cm1479 & Cm2683) in the 25S rRNA, are plant-specific, since equivalent positions are also found in the rRNA sequences in tomato (57) and tobacco (58) but not in the human or yeast rRNA sequences (reviewed in (34)).

The Arabidopsis genome encodes a total of 226 C/D-box snoRNA genes, 143 of which target the 117 rRNA 2’-*O*-methylation sites (12). We investigated potential changes in the accumulation of C/D-box snoRNAs targeting the 12 identified Nm sites. Differentially expressed C/D-box snoRNA (logFC ≥ 2 or logFC ≤ −2) include 45 in DS v/s 4d, 26 in DS v/s 15d and 1 in 4d v/s 15d (**Fig. 5c**). Only two of these C/D-box snoRNAs (At2gCDbox68.1 and At5gbox136.1,) target Nm sites with 2’-*O*-Me changes during seed-to-seedling transition (**Fig. 5c**). In the DS v/s 4d comparison, At5gC/Dbox136.1 and 25S_Cm2683 showed a positive correlation, while At2gCDbox68.1 and 5.8S_Gm79 exhibited a negative correlation.

Taken together, these results suggest that 2’-*O*-Me changes of specific Nm is dynamic during Arabidopsis seedling establishment. Additionally, the data indicate that the expression of C/D-box snoRNAs is not fully correlated with 2’-*O*-Me levels at their respective target sites.

### Polysome profiling reveals dynamic and transcript-specific ribosome loading during the seed-to-seedling transition

Changes in 2′-*O*-Me levels at specific Nm sites between DS, 4-day-old seedlings, and 15-day-old seedlings prompted us to investigate whether dynamic rRNA methylation affects translational activity during the transition from seed to stablished seedlings. For this purpose, we performed polysome profile and RNA-seq on DS, DS sown on MS medium for 48 h at 4°C in the dark (SS), 4- and 15-day-old seedlings.

Polysome profiling revealed higher accumulation in 4-day-old seedlings than in 15-day-old seedlings, with the lowest accumulation observed in SS. While the 80S and 40S/60S subunits, as well as polysome peaks, were detectable in 4- and 15-day-old seedlings, SS samples exhibited a single, broad increasing peak, lacking resolved polysome profiles. No 80S, 40S/60S subunits, or polysome profiles could be obtained from DS. Consistent with previously published profiles (47), polysome accumulation was greater in 4-day-old seedlings than in 15-day-old seedlings (**Fig. 6a**), suggesting that rRNA methylation dynamics may directly or indirectly affects polysome-associated transcript accumulation in during seed-to-seedling transition.

**Figure 6:**
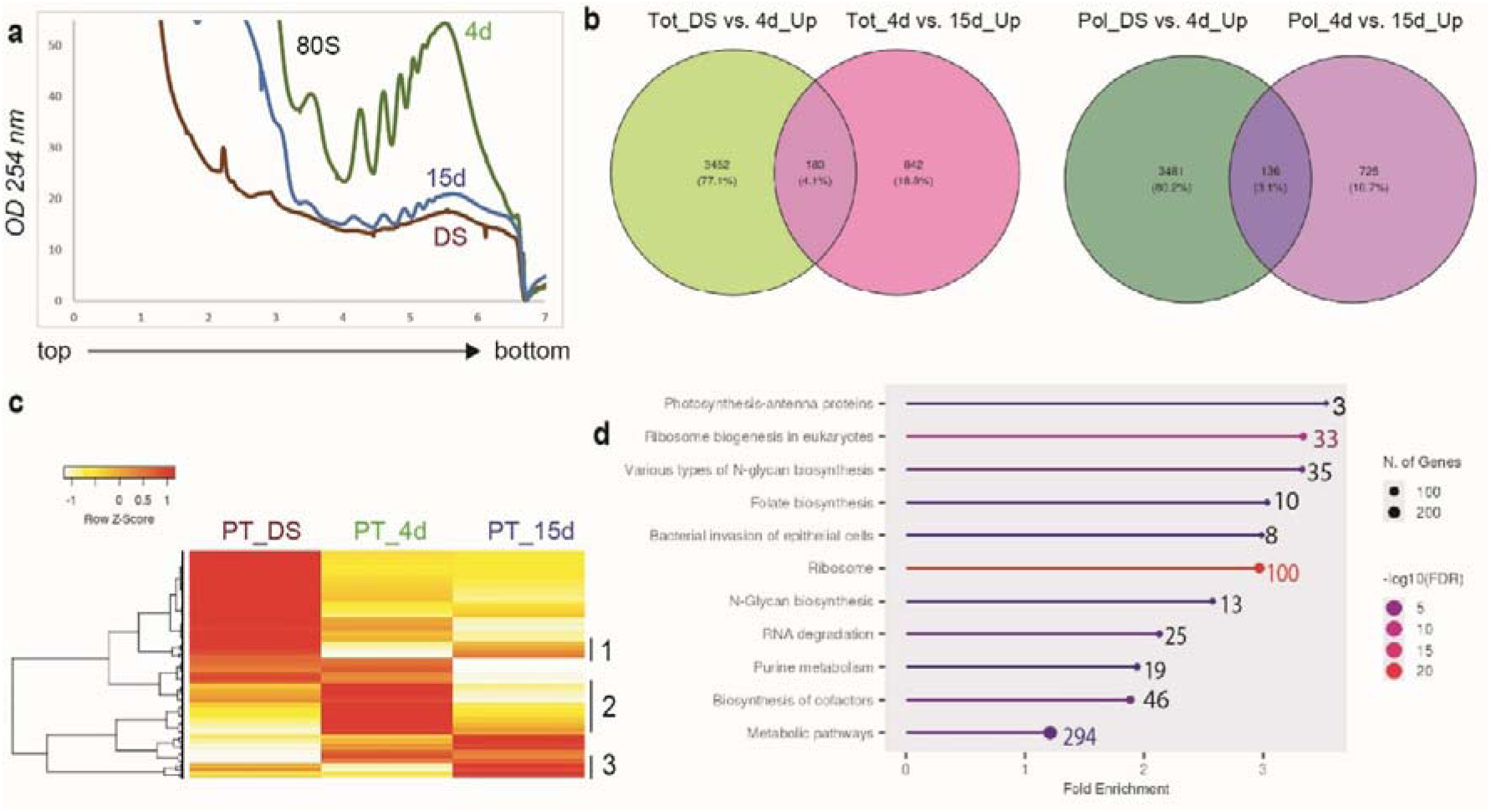
Polysome profiling and transcript-specific loading during seed-to-seedling transition. **a**, Polysome profiles (replicate 1) from 48 h stratified seeds (SS, brown), 4-day-old (green) and 15-day-old (blue) seedlings. **b,** Venn diagrams show the distribution of significantly upregulated (FC ≥ 1.5, p-value ≤ 0.05) mRNA across SS vs. 4d and 4d vs. 15d seedlings in total and polysome RNA. **c,** Heat map of transcripts ratio (PT = RPM in Polysomes / RPM in total RNA) in SS, 4-day-old seedlings, and 15-day-old seedlings. Cluster 2 (C2) shows accumulation of transcripts while clusters 1 (C1) and 3 (C3) show decreased accumulation of transcripts in 4-day-oldseedlings **d)** Fold-enrichment analysis (using ShinyGO 0.85.1, https://bioinformatics.sdstate.edu/go/) of polysome-associated RNA transcripts accumulating (FC ≥ 1.5, *p*-value ≤ 0.05) in 4-day-old seedlings. Numbers indicate the count number of gene transcripts per gene category.

To determine this impact, RNA-seq analysis was performed on total and polysome-associated RNAs from SS, 4-, and 15-day-old seedlings. PCA demonstrated that biological replicates for each condition clustered tightly, with distinct separation between SS, 4-day, and 15-day samples. The total RNA-seq data identified 6,776 (3,635 up and 3,141 down) differentially expressed genes (DEGs) in SS vs. 4-day-old seedlings, and 1,875 DEGs (1,025 up and 850 down) in 4-day-old seedlings vs. 15-day-old seedlings. In polysome fractions, 7,303 RNA sequences with differential polysome association (3,617 higher and 3,686 lower loading) were detected in SS vs. 4-day-old seedlings and 1,413 (861 higher and 552 lower loading) in 4-day-old seedlings vs. 15-day-old seedlings (**Fig. 6b**). Among these, 424 and 170 transcripts with higher polysome loading were not differentially expressed in SS vs. 4-day-old seedlings or 4-day-old seedlings vs. 15-day-old, respectively.

For each transcript, polysome accumulation was assessed by comparing its RPM in the polysomal fraction with its RPM in the total RNA fraction. The dynamics of the resulting ratio across seedlings establishment was assessed by a clustering analysis using hierarchical heatmap. Interestingly, three distinct clusters of transcripts that correlates with ribosomal 2’-*O*-Me levels dynamic were identified (**Fig. 6c**). Cluster 2 included 2,872 transcripts with enhanced polysome loading in 4-day-old seedlings compared to the two other stages, primarily encoding proteins involved in photosynthesis and ribosome biogenesis. Clusters 1 and 3 comprised 907 and 796 transcripts, respectively, characterised by reduced polysome loading in 4-day-old seedlings but elevated loading in SS and 15-day-old seedlings. Among the polysome-enriched transcripts (≥1.5-fold), the most represented functional categories included: ribosomal proteins, ribosome biogenesis factors, cofactor biosynthesis, N-glycan biosynthesis, RNA degradation pathways (**Fig. 6d**). We searched for conserved RNA sequence motifs in the 5’ and 3’ untranslated regions (5’ and 3’ UTRs) of the 2,907 transcripts in cluster 2. However, this analysis did not reveal any significantly enriched motifs associated with preferential polysome loading at four days.

Collectively, these data suggest that dynamic changes in rRNA methylation during seed-to-seedling transition may contribute, directly or indirectly, to the modulation of translation, thereby helping fine-tune the translational program.

## DISCUSSION

To better understand the regulation and functional impact of rRNA 2’-*O*-Me, we characterised *Arabidopsis nuc1-2* plants, which display growth and developmental defects and exhibit constitutively reduced rRNA 2’-*O*-methylation levels (12), and investigated its dynamics during seed-to-seedling transition (see **Fig. 7**).

**Fig. 7:**
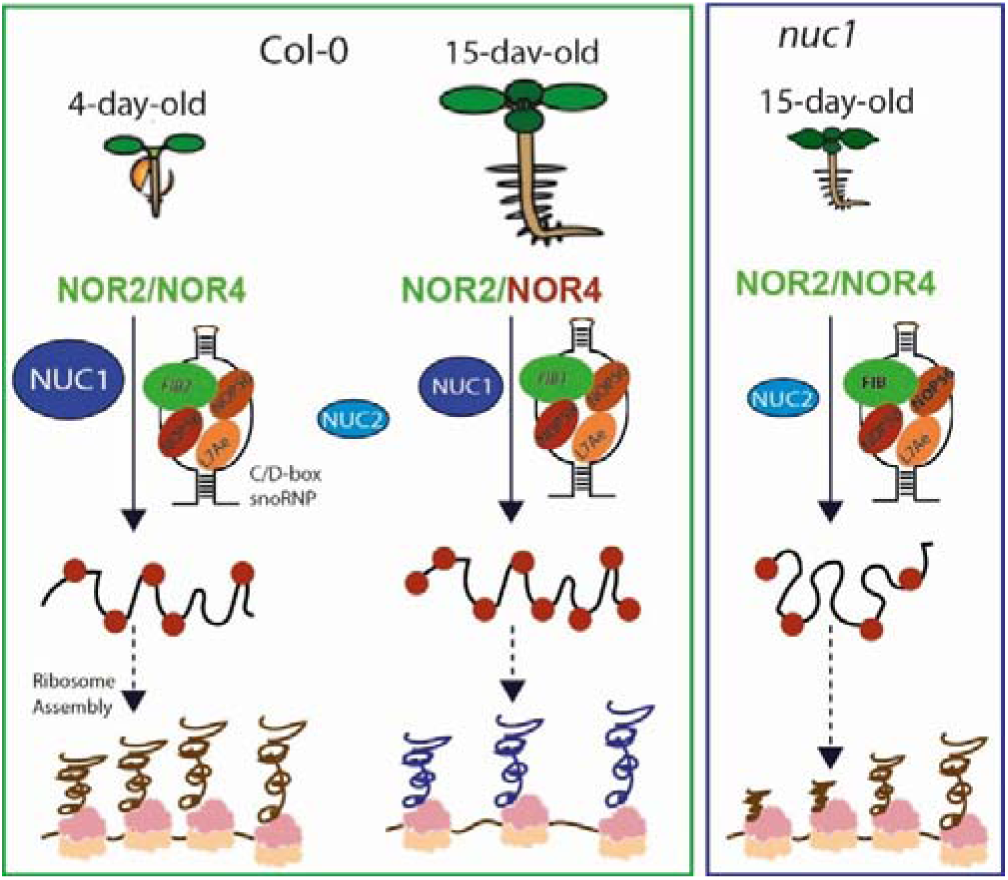
Global overview. In 4-day-old seedlings and *nuc1-2* plants, rRNA genes are expressed from NOR2 and NOR4, and rRNAs are hypomethylated. The observed changes in 2′-*O*-Me appear unrelated to NORs expression, the differential expression of C/D-box snoRNPs, or the NUC2 protein, which is either transiently expressed during seedling establishment or constitutively expressed in the absence of NUC1. Hypomethylated rRNAs, to varying extents in both 4-day-old seedlings and *nuc1-2* plants, lead to polysome accumulation and affect the translation of specific mRNAs expressed in *nuc1-2* and during seedling establishment, including transcripts encoding ribosome related protein (brow nascent polypeptides).

Changes in 2’-*O*-Me levels at specific Nm sites, observed in *nuc1-2* mutants and during seed-to-seedling transition appear largely independent of differential C/D-box snoRNA expression. Consistently, snoRNA expression levels also fail to account for the differential rRNA modification levels across mammalian tissues (Milenkovic et al., 2025). Nevertheless, C/D-box snoRNAs that are differentially expressed may be involved in the 2′-*O*-Me of non-rRNA targets. C/D-box snoRNAs are known to guide 2′-*O*-Me not only in rRNAs but also in tRNAs, snRNAs, snoRNAs, lncRNAs, and mRNAs, thereby influencing translation through diverse regulatory mechanisms (reviewed in (59–61)).

Furthermore, in *nuc1-2* plants and during seed-to-seedling transition, all C/D-box snoRNP core proteins and snoRNAs are expressed, co-purify with fibrillarin and localise in the nucleolus in *nuc1-2* plants, indicating that C/D-box snoRNP assembly is likely unaffected (**Figs. 1-3**). Surprisingly, fibrillarin proteins accumulate in *nuc1-2* plants, contrasting with previous studies in cancer cells where FIB upregulation increases rRNA 2′-*O*-Me (24,25). However, fibrillarin accumulation is unlikely to inhibit 2’-*O*-Me, as overexpression of FIB2–YFP in Col-0 plants does not affect rRNA 2’-*O*-Me.

During seed-to-seedling transition and in *nuc1-2* plants both Nucleolus Organiser Regions (NOR2 and 4) are active and express specific rRNA gene variants (**Fig. 7** and (40,41,62)). However, analysis of Arabidopsis accessions Can-0 (Canary Islands) and Bur-0 (Burren), which express NOR2 and NOR4, respectively (63), neither *nuc2-2* with actives NOR2 and 4, failed to establish a correlation between changes in 2’-*O*-Me and the expression of NORs.

Transcription, processing of pre-rRNA and 2’-*O*-Me of rRNA occur mostly in the nucleolus (4,17,64,65). In *nuc1* mutants, the functional nucleolar structure is compromised, with disappearance of FCs and strong reduction of the DFC (37). The marked reduction of DFC in *nuc1* mutants, may inhibit 2′-*O*-Me reactions, which normally occur co-transcriptionally within this compartment. Indeed, 2’-*O*-Me remains unaffected in *apum23* plant mutants despite their compromised pre-rRNA processing. Yet, it remains unclear how this disorganisation selectively affects specific rRNA sites rather than all of them. Further studies are therefore required to determine the impact of pre-rRNA structure on the accessibility of C/D-box snoRNPs at specific Nm sites and, consequently, 2’-*O*-Me activity.

The accumulation of polysomes in *nuc1-2* plants are unlikely to result directly from the loss of NUC1 itself, particularly given that NUC1 does not associate with polysomes. Our findings align with studies in mice, where nucleolin (NCL) interacts with the free mRNA and 40S fractions but not to polysomes (66). The incorporation of incompletely processed pre-rRNAs into ribosomes may also inhibit or prevent efficient translation (67,68). However, we exclude this possibility, as 5’ and 3’ extremities of rRNAs incorporated into ribosomes are accurately processed in *nuc1-2* plants.

A recent study shows that, reducing ribosome load improves mRNA translation in human cells (69). Therefore, in *nuc1-2* plants, the accumulation of polysomes with almost unchanged protein expression profiles suggests that highly loaded polysomes may compensate for modest ribosome translation initiation/elongation rates (**Figs. 5**). Indeed, the 5’Pseq analysis revealed an accumulation of 5’P reads at −13nt relative to the ATG start codon, which corresponds to the position of a ribosome stalled at the P site during the initiation step, as described in yeast cells (48). Furthermore, additional pauses were observed at the stop and other codons, suggesting that reduced 2’-*O*-Me at specific rRNA sites impact different aspects of translation in *nuc1-2* plants. The arrest or inhibition of translational elongation from the ATG start or other codon is unlikely to be due to a lack of tRNA, as no major differential expression is observed in *nuc1-2* mutants. In this context, increased loading of mRNA encoding ribosome-related proteins can be interpreted as feedback signalling directed to increase and compensate the stalled ribosome activity in *nuc1-2*.

Differentially methylated Nm sites during seed-to-seedling transition are also demethylated in *nuc1-2* mutants and are located in the different domains of the rRNAs (**Fig. 5**), including the 25S domain V, which is responsible for the peptidyl-transferase activity and tRNA binding (70), suggesting that these rRNA 2’-*O*-Me changes could be required for tuning translation activity. Specific rRNA modifications may contribute to the assembly of heterogenous and specialized ribosomes, enabling the translation of particular mRNAs during plant development (71).

Notably, reduced 2’-*O*-Me of specific rRNA sites is also accompanied with an increased polysome accumulation and differential mRNA-polysome loading (**Fig. 6**). The enrichment of polysomes with transcripts encoding ribosomal proteins, ribosome biogenesis factors, and rRNA expression from both NORs strongly indicates a privileged enhancement of ribosome biogenesis. Consistently, during seedling establishment, plants must synthesize ribosomes and selectively translate mRNAs to transition from reliance on seed reserves to autonomous growth. These low 2′-*O*-Me ribosomes may further optimize the translational efficiency of mRNAs involved in photosynthesis-related proteins and glycogen biosynthesis pathways, both of which are essential for successful seedling establishment in Arabidopsis (72–74).

Few or non-polysomes were also detected in dormant dry seeds, while similar polysome accumulation occurred in early imbibed dormant seeds and imbibed non-dormant seeds under dark and 25°C conditions (75). Under light at 22°C., polysome accumulation was also observed in imbibed non-dormant seeds (76). Recently, dormancy release and early imbibition were also shown to induce gene-specific changes in translation efficiency, independent of transcript abundance (77). Profiling 2’-*O*-Me during these stages and conditions should help us to better understand the role of ribosome modifications in plant development.

We have not yet identified specific RNA sequences that could explain the polysome loading profiles observed during seedling establishment. In human cancer cells, hypermethylated (2’-*O*-Me) ribosomes translate mRNA with reduced fidelity and increase internal ribosome entry site (IRES)-dependent translation initiation of key cancer genes (24,25). To date, WUSCHEL mRNA is the only known mRNA in Arabidopsis, that initiates translation via an IRES-dependent mechanism. Notably, the AtLA1 protein binds to the 5′-untranslated region (5’UTR) of WUSCHEL and initiates the IRES-dependent translation (78). In this context, the accumulation of AtLA1 protein, observed in *nuc1-2* plants or the differential expression of nucleolin proteins during seedling establishment (41), may be required to regulate translation of specific mRNAs. It remains unknown whether plant nucleolin interacts with 5′ or 3′UTR sequences and/or regulates translation. However, mammalian nucleolin has been implicated in the translation via its interaction with internal ribosome entry site (IRES)-containing mRNAs (66) or via G-rich elements in the 5’ or 3’UTR sequences (61). Similarly, C/D-box snoRNAs that differentially accumulate in *nuc1* and/or during seed-to-seedling transition might also affect mRNA loading and/or translation in the hypomethylated ribosomes. Indeed, snoRNA can affect translation either through direct mRNA interaction or mRNA methylation (61).

The alterations in mRNA loading and translation activity may also arise from disruptions in the assembly of the translation initiation complex, global ribosome structure, or specific functional domains, such as the peptidyl transferase center (PTC) or the decoding center. Further studies are required to elucidate how 2’-*O*-Me regulation of specific rRNA sites modulates ribosome activity in plants. A recent study reported that loss of 2′-*O*-Me in yeast ribosomes impairs the translation of ribosomal protein transcripts by altering codon usage and ribosomal structure (79). Therefore, structural and computational modelling studies are essential to address this issue in plants, particularly to investigate both global and specific 2′-*O*-Me modifications during development.

## METHODS

### Plant materials and growth conditions

*Arabidopsis thaliana* Col-0, Can-0, and Bur-0 were obtained from the Nottingham Arabidopsis Stock Centre (http://nasc.life.nott.ac.uk). The *nuc1-2* (SALK_002764), *nuc2-2* (GABI_178D01), *apum23-1* (SAIL_757_B08 and *apum23-2* SALK_052992), plant mutants were described in (37,40,41,46). The *fib1* (GK-322B02) mutant contains a T-DNA insertion in the 5^th^ exon of the FIB1 gene. The expression of the chimeric FIB2-YFP protein is under the control of the cauliflower mosaic virus 35S RNA (35S CaMV). Col-0 and *nuc1-2* plants expressing FIB2-YFP were previously described in (12,40–42). Seeds of the 35S::FH-RPL18 plant line described in (44) were obtained from the Arabidopsis Biological Resource Center (ABRC). To obtain plants expressing EGFP-NUC1 and NUC1-EGFP chimeric proteins, Agrobacterium-mediated transformation of *nuc1-2* plants was performed with a pK7WGF2 (Gateway) plasmid containing EGFP (Enhanced Green Fluorescence Protein) and NUC1 coding sequences under the control of the 35S CaMV promoter sequence.

Seeds were sown either on soil (4/5 soil and 1/5 vermiculite) or on 1X Murashige and Skoog (MS) medium (MS containing 1% sucrose). After 2 days of stratification at 4°C in the dark, plants on soil were grown for 3 weeks in controlled growth chambers under a 16-hour light/8-hour dark cycle at 20°C with a light intensity of 100 µE.m-2.s-1 and a relative humidity of 60% (CLF Plant Climatics GmbH, Wertingen, Germany) while plants on MS medium were grown under continuous light (100 µE.m-2.s-1) for 4-15 days. Aerial parts (soil grown) or whole seedlings (grown *in vitro*) were collected, shock frozen in liquid nitrogen, ground to a fine powder, and stored at −80°C. Herein, a biological replicate always consists of several individual plants from the same line, at the same age, grown under the same conditions and in the same incubator at the same time.

### RNA isolation

To obtain total RNA, approximately 100 mg of dry seeds (DS), stratified seeds (SS), or seedlings, were first frozen in liquid nitrogen and then ground into a powder using a mortar and pestle. For RiboMethSeq, RNA-seq, Mim-tRNA-seq, and 5’PSeq analysis, RNA was extracted using the Monarch® Total RNA Miniprep Kit (NEB), according to the manufacturer’s instructions, including optional DNase treatment. For semiquantitative RT-PCR total RNA was extracted using TRIzol™ as previously described (12). The RNA was then precipitated with >95% ethanol and resuspended in 50 µL of nuclease-free water. Total RNA from stratified seeds was extracted using the Monarch® Total RNA Miniprep and the RNAqueous (Invitogen) kits.

To obtain total RNA from polysomes, 250 µL of each fraction was mixed with 750 µL Trizol and incubated for 5 min at RT. Then 200 µl of cold chloroform was added and the samples were incubated for 2-3 min at RT, and the mixture was centrifuged at 8,000 × g for 15 min at 4°C. The aqueous phase was collected and precipitated with 200 µl of isopropanol. After 10 min incubation at RT, the mixture was centrifuged at 8,000 × g for 10 min at 4°C. The isopropanol was removed, and 1 mL of 70% ethanol was added to the pellet (and mixed 6-7 times by inverting the tube). The mixture was centrifuged at 8,000 × g for 3 min at 4°C, the ethanol was removed, and the pellet was air dried under the hood for 5 min. The RNA was dissolved in 50 µL of nuclease-free water.

The quantity and quality of the RNA were verified using the Agilent RNA 6000 Nano Kit (Agilent, 5067-1511) and analysed on a Bioanalyzer 2100 expert_Plant_RNA nano with Agilent RNA 6000, according to the manufacturer’s instructions.

### RT-PCR

First-strand cDNA synthesis was performed on 1.5 µg of DNase-treated RNA using the random and oligo dT primers together with the GoScript Reverse Transcription System kit according to the manufacturer’s instructions (Promega, A5003). PCR amplification of FIB1 and 3’ETS sequences was performed using a PTC-200 device (MJ Research) and oligo pairs o1279/1282 and *o108/0109*, respectively. Oligo pairs *5ef1a/3ef1a* were used to amplify eIF1α transcripts, which were used as an internal loading control.

### Polysome profiling

Polysome profiling was performed on the aerial parts of 18-day-old Col-0 and *nuc1-2* plants grown on soil, as well as from dry seeds sown on MS medium for 48 h at 4°C in the dark and from *in vitro*-grown seedlings 4- and 15 days old, as described in (80) with minor modifications. The harvested plant seedlings were frozen in liquid nitrogen and cryogenically ground using the Retsch MM400 cryobroyer. The plant seedling powder (400 mg) was suspended in 1.2 mL of polysome extraction buffer (200 mM Tris•HCl pH 9; 200 mM KCl; 25 mM EGTA; 35 mM MgCl2; 1% Detergent Mix (1% [w/v] Brij-35; 1% [v/v] Triton-X100; 1% [v/v] Igepal; 1% [v/v] Tween-20; 1% [w/v] Sodium Deoxycholate; 0.5% [w/v] Polyoxyethylene (10) tridecyl ether); supplemented with 5 mM dithiothreitol; 50 µg/mL Cycloheximide; 50 µg/mL Chloramphenicol; 1x MG132 protease inhibitor (Calbiochem; Cat: 474790-20MG) and 1% [v/v] protease inhibitor cocktail [Sigma-Aldrich]. Then, 900 µL of the mixture was resolved on a 15–60% sucrose gradient (9 mL) and centrifuged at 209,678 × g for 3.5 h (rotor SW41-Ti, Beckman Coulter). Polysome profile analyses were monitored at 254 nm. Sucrose gradient fractions (600 µL) were collected using a BioLogic Duoflow fraction collector (Bio-Rad Laboratories). To obtain fraction-wise protein samples throughout the polysome gradient, 12 fractions were collected. To obtain total RNA from 40S/60S and polysomes, respectively, peak-wise collection was performed taking into account the time delay between the detector and the fractionator. Upon detection of the 40S/60S or first polysome peak, respectively, a timer was set to 40s, after which the collection was switched to a new tube. The same was done to define the end of the 40S/60S pick and last polysome peak, respectively. The resulting peak-specific fractions corresponding to the 40S/60S and polysome fractions, respectively, and were pooled prior to RNA extraction.

For protein precipitation of the polysome fractions, two volumes of 100% cold ethanol were added to each sample and incubated for 6 h at 4°C. The precipitates were then centrifuged for 30 min at 4°C and 16,000 × g, washed twice with 1 mL of 70% cold ethanol, dried and resuspended in 4× Laemmli buffer (input in 50 µL, fraction 1-4 100 µL and fraction 5-12 in 10 µL). For immunodetection, 5 µL of each input fraction and 10 µL of each polysome fraction were treated as described in the section “Protein extraction and Western Blot analysis”. Membranes were hybridised overnight at 4°C with either α-GFP (1:5,000, Rockland), α-RPL13 (1:10,000, Agrisera AS13 2650) or α-RPS10 (1:20,000: previously reported in (6)).

### RiboMethSeq

The RiboMethSeq analysis was performed as described in (12), using the herein validated Nm positions present in *A. thalian*a rRNA (12,33). Briefly, 100 ng of total RNA was subjected to alkaline hydrolysis at 96°C for 12 min. The precipitated and end-repaired RNA was then converted into a library using the NEBNext© Small RNA Library Kit (NEB, USA). The quality and quantity of the library was assessed using a High Sensitivity DNA Chip on a Bioanalyzer 2100 and a Qubit 2.0 fluorometer. High-throughput sequencing of the multiplexed libraries was performed on an Illumina NextSeq 2000 instrument in a 50 nt single-end read mode. RiboMethSeq analysis measures the protection of a given phosphodiester bond against alkaline hydrolysis. Thus, the calculated scores reflect the level of protection, which correlates with 2’-*O*-methylation level. If a nucleotide is not protected against cleavage or even more intensively cleaved compared to the neighbouring nucleotides, the calculated score is close to zero or even negative. This is typically observed for very low modified or unmodified positions. Normalisation of the raw RiboMethSeq scores was performed to the average value by position, observed in whole treated series of samples (81–83). Detailed information on the methodology of RiboMethSeq methodology and analysis is provided in Supplementary Information.

### RNA-seq of C/D-box snoRNA and bioinformatic analysis

For C/D-box snoRNA expression analysis we used total RNA from aerial parts (18-day-old Col-0 and *nuc1-2* plants), as well as from dry seeds and *in vitro-*grown seedlings 4 and 15 days old. Total RNA was extracted using the Monarch® Total RNA Miniprep Kit (NEB) from biological triplicates. 50 ng of total RNA was directly converted into a library using the NEBNext© Small RNA Library Kit (NEB, UK). The quality and quantity of the library was assessed using a High Sensitivity DNA Chip on a Bioanalyzer 2100 and a Qubit 2.0 fluorometer. High throughput sequencing of the multiplexed libraries was performed on an Illumina NextSeq 2000 instrument in a 50 nt single end read mode.

Reads were cleaned with trimgalore (http://www.bioinformatics.babraham.ac.uk/projects/trim_galore/), keeping only reads longer than 40 nt. Cleaned reads were mapped with bwa mem (84) onto the *A. thaliana* snoRNAs and snRNAs. The gene read count was performed with featureCounts (85). We used the IDEAmex (Integrated Differential Expression Analysis MultiEXperiment) web application (86) to perform data exploration and differential expression analysis. The input of the IDEAmex pipeline was the raw count table generated from the previous step for all replicates and conditions. PCA (Principal component analysis) plots were obtained from normalised data. Differential expression analysis was performed in IDEAmex by using DESeq2 (87), limma-voom (88), EdgeR (89) and NOISeq (90) methods. To identify differentially expressed genes, we used default parameters (CPM ≥ 1, absolute LogFC ≥ 1, adjusted *p*-value or FDR ≤ 0.05). We checked the TOP box to keep genes differentially expressed between EdgeR, Limma, DESeq2 and NOISeq. The MA plot showing the logarithm of the fold-change (logFC) and the average log count (logCPM) was generated using the ggplot2 package and, as inputs, the tables that contain respectively the EdgeR output (S5.2) and the IDs of C/D-box snoRNA that are at the intersection of the DEG obtained by all the different selected methods (S5.3).

### RNA-seq and 5’PSeq

For transcriptomic analysis, total and polysomal RNA were prepared from Col-0 and *nuc1-2* mutant plants (biological triplicates), as well as from dry seeds sown on MS medium for 48 h at 4°C in the dark and from *in vitro-*grown seedlings 4- and 15 days old. Library preparation and sequencing was performed by Azenta/GENEWIZ, from strand-specific RNA-seq using polyA selection libraries and Illumina NovaS-seq to generate 2×150 bp-long reads.

For 5’PSeq (5′ monophosphorylated mRNA degradation intermediates sequencing) analysis, RNA was extracted from 3 biological replicates of aerial parts from 18-day-old Col-0 and *nuc1-2* plants. Library preparation was performed as previously described (47).

### Bioinformatic analysis of mRNA-seq and 5’PSeq

For RNA-seq data, adapters and low-quality reads were removed. For each library, reads were trimmed using Trimmomatic v0.39 (91). Trimmed reads were filtered out from reads corresponding to chloroplastic, mitochondrial, ribosomal and small RNA sequences using bowtie2 v2.4.4 (92) in sensitive-local mode. Reads were then mapped against TAIR10 genome and the gtf Araport11 annotation file using Hisat2 v2.2.1 (93) with standard parameters. Only unique mapped reads were kept using samtools v1.13 with option ‘-q 10’ (94). Reads count by gene was performed by htseq-count in union v1.99.2 in ‘union’ mode (95). Differential expression analyses were performed using the Bioconductor R package DESeq2 (87), with a false discovery rate of 0.05. P-values were corrected for multiple tests by the Benjamin-Hochberg rule (adjusted *p*-value).

For 5’Pseq data, as sequencing was performed in paired-end, only reads corresponding to R1 were analysed and trimmed to 50 nt using Trimmomatic v0.39 (91). Reads were then mapped as described for RNA-seq data. Metagene analysis was performed by the FivePSeq software (50). ‘meta_counts_START.txt’ and ‘codon_pauses.txt” files were used to analyse 5′P reads accumulation around start and codon-specific positions.

Metagene analysis of transcripts significantly affected in *nuc1* was performed with FIVEPSEQ by rerunning the analysis on the corresponding transcript subset using the same parameters as for the whole dataset.

To identify sequence motifs in 5’UTRs and 3’UTRs, Araport11 annotations are used and analysed using HOMER (https://bio.tools/homer)

### Mim-tRNA-seq

For Mim-tRNA-seq, total RNA was extracted from aerial parts of 18-day-old Col-0 and *nuc1-2* plants grown on soil and from biological triplicates. The total RNA was then subjected to an additional LiCl precipitation step that allows enrichment, in the supernatant, of RNAs of a size smaller than 150 nt. Briefly, 200 µL of a LiCl solution (5 M LiCl, 50 mM Tris•HCl pH 7.5) was added to 300 µL of RNA (0.5 µg/µL) and the mixture was incubated at 4°C on ice for 4 hours. Then, the rRNAs and mRNAs were pelleted by 30 min centrifugation at 13,000 × g. The aqueous phase was collected and subjected to an ethanol precipitation (96). The tRNA fractions were sequenced using an adaptation of the mim-tRNA-seq protocol (97). Biological triplicates from WT and *nuc1-2* mutant plants were analyzed. Data analysis was performed using the dedicated bioinformatics pipeline developed in (98). Only reads longer than 10 bp with quality >30 were kept for the mim-tRNA-seq analysis. The package mim-tRNA-seq version 1.3.8 (https://github.com/nedialkova-lab/mim-tRNA-seq) was used with the following parameters: --cluster-id 0.90 --min-cov 0.0005 --max-mismatches 0.1 --max-multi 4 --remap --remap-mismatches 0.075 --control-condition total WT. Sequences were mapped to annotated tRNAs of *A. thaliana* extracted from PlantRNA (http://plantrna.ibmp.cnrs.fr/) and reformated for mimseq. The bam files were sorted and indexed with samtools version 1.16 (99). Differential analysis of tRNA expression was performed with the DESeq2 package version 1.42.0 from R-Bioconductor (87).

### Immunoprecipitation (IP): Protein and RNA extraction

Leaves from 21-day-old Col-0+FIB2-YFP and *nuc1-2*+FIB2-YFP plants (5 biological replicates) were ground to a fine powder in liquid nitrogen. Whole cell extracts were prepared in 3 volumes of EB150 buffer (50 mM Tris•HCl pH 7.5, 150 mM NaCl, 5 mM MgCl2, 10% glycerol, and EDTA-free protease inhibitor cocktail [Roche, 11836170001]) supplemented with 0.1% NP-40. Cell extracts were incubated with GFP-Trap_MA beads (Chromotek) for 2 h at 4°C with gentle rotation. The unbound fractions were collected, and the beads were washed 3 times with EB150. Bead-bound proteins were recovered using 4× SDS-Laemmli buffer (250 mM Tris•HCl pH 6.8; 8% SDS; 40% glycerol; 0.2% bromophenol blue, 0.4 M DDT) for Western blot analysis and for LC-MS/MS analysis as described in (12).

For RNA extraction, RNA from the IP fraction was extracted using TRI Reagent (MRC research), as described in (12). RNA separated in aqueous phase were precipitated using glycogen as a carrier (SIGMA, 20 mg) following supplier instructions. The quantity and quality of the RNA were verified as described in section RNA isolation.

### Fluorescence in situ hybridisation (FISH)

RNA-FISH was performed on *A. thaliana* roots from 7-day-old seedlings grown on MS medium, according to the protocol described in (60). The sequences of the 3’-end labelled cyanine (Merck) antisense oligonucleotides U3, At5gCDbox137.1/snoR29, At1gCDbox25.1/HID2, At1gCDbox33.1, At5gCDbox141.1, At4gCDbox117.1 and At1gCDbox18.1 snoRNAs are given in **Table 1**.

**Table 1:**
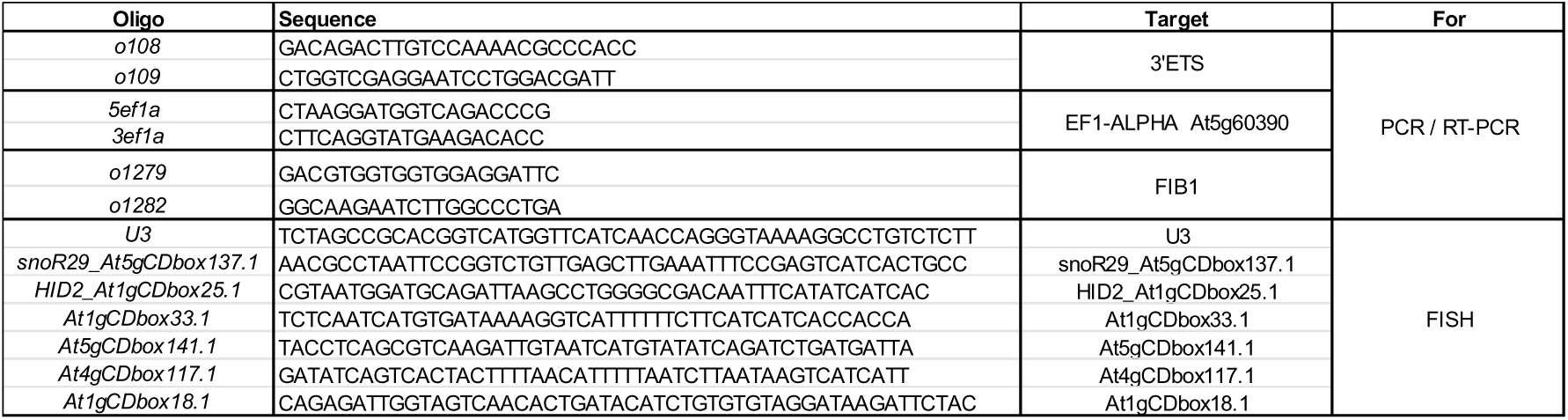
List of primers.

### Immunocytofluorescence

Immunofluorescence was performed on roots apex from 8-day-old (Col-0 and *nuc1-2*) seedlings grown on MS medium without sucrose. Whole seedlings were fixed for 20 min with 4% formaldehyde in cold Tris buffer (10 mM Tris•HCl pH 7.5, 10 mM EDTA, 100 mM NaCl) and washed twice for 10 min with Tris buffer. The root apices were dissected, placed on a microscope slide and treated with digestion solution (1% cellulose, 0.5% cytohelicase, 1% pectolyase in Tris buffer) during 2 h at 37°C and washed twice with Tris buffer. The root apices were squashed on polysine slide and frozen in liquid nitrogen. The slides were treated with solution 1× PBS (137 mM NaCl, 2.7 mM KCl, 10 mM Na_2_HPO_4_, 2 mM KH_2_PO_4_ pH 7.5) supplemented with 0.5% Triton-X100, for 15 min at room temperature and then washed twice for 5 min with 1× PBS. Slides were incubated with 120 µl of hybridisation solution (1× PBS supplemented with 3% BSA and 0.005% Tween-20) with α-NOP56 (1:500, GenScript) or α-Fib (1:500, 38F3 Abcam) at 4°C overnight. Slides were washed twice for 5 min with 1× PBS and hybridised with Alexa 488 (1:1000, Invitrogen) in 1× PBS (supplemented with 3% BSA and 0.005% Tween-20) for 2 h at room temperature. Slides were then washed twice for 5 min with 1× PBS and mounted in Vectashield medium containing 4′,6-diamidino-2-phenylindole (DAPI) solution 1 µg/ml. Observations and imaging were performed using a confocal microscope LSM 900 Airyscan 2 from Zeiss and images treated with Fiji programme.

### Protein extraction and Western blot analysis

Total proteins were extracted from 21-day-old Col-0, *nuc1-2*, 35S::FIB2-YFP (in Col-0 or *nuc1-2* backgrounds), and *nuc1-2* complemented 35S::EGFP-NUC1 or 35S::NUC1-EGFP plants. 100 mg of leaves was ground to a fine powder with liquid nitrogen and then resuspended in 300 µL of 2× Laemmli buffer (125 mM Tris•HCl pH 6.8; 4% SDS; 20% glycerol; 0.1% bromophenol blue) supplemented with β-Mercaptoethanol (0.5% final). The mixture was incubated at 65°C for 10 min, centrifuged at 14,000 × g for 15 min and the supernatant was retained as the total protein extract.

For Western blot analysis, 15 µL of total protein extract or immunoprecipitated fractions were subjected to 10% SDS PAGE, transferred to PVDF (Millipore) or nitrocellulose (BioRad) membranes, followed by Western blot analysis as previously described (12). Membranes were hybridised overnight at 4°C with either α-GFP (rabbit polyclonal GFP antibody with reactivity against GFP and YFP according to manufacturer’s information, 1:5,000, Rockland), α-FLAG-HRP (1:10,000, Sigma), α-HA-HRP (1:12,000, SIGMA), α-FIB 38F3 (1:250, Abcam), α-TUBULIN (1:10,000, Agrisera), α-NOP56 :(1:1000, GenScript), α-UGPase (1:10000, Agrisera), α-H3 CT pan (1:10000, Merck Millipore) or α-ACTINE (1:5000, Life Technology). Antibodies α-NOP56 were custom raised against the peptide sequence NLPKVKEGKKPKFSC, selected from the NOP56 gene At1g56110 (GenScript). α-NUC1 (1:100,000) and α-RPL13 (1:5,000) and α-RPS6 (1:5,000) were previously reported in (40,100,101). Immunoreactive proteins were detected using Immobilon western chemiluminescent substrates (Millipore, P90720). The images were acquired using the Fusion Solo S camera (Vilber Lourmat). After immunodetection, the PVDF membranes were stained with colloidal blue Coomassie solution, and the nitrocellulose membranes were stained with Ponceau Red solution.

### Liquid Chromatography-Tandem Mass Spectrometry (LC-MS/MS) analyses

Total protein extracts (5 biological replicas) and the IP eluate protein fractions (biological triplicates) from Col-0+35S:FIB2-YFP and *nuc1-2*+35S:FIB2-YFP were diluted to a 1X SDS-Laemmli buffer and supplemented with 10 mM DTT. The mixture was then loaded onto an in-house cast 4% acrylamide stacking gel. After staining the gel with Coomassie Blue, the stacking bands were manually excised and subjected to nanoLC MS/MS analysis as described in (12). Detailed information on the nanoLC MS/MS methodology is provided in Supplementary Information.

## Data availability

The RNA-seq data have been submitted to the Sequence Read Archive (SRA): Total RNA (PRJEB36061), small RNA (PRJNA1097563), nuc1 polysome RNA (PRJNA1106377) and 5’PSeq (PRJNA11007736), IP-FIB2 small RNA (PRJNA1477268) and seedling establishment polysome RNA PRJNA1498475. The RiboMethSeq data have been submitted to the <u>ENA Browser (ebi.ac.uk)</u> with the identifier PRJEB75969 and PRJEB115598. https://www.ebi.ac.uk/ena/browser/home The mass spec-trometry proteomics data have been deposited in the ProteomeXchange Consortium via the PRIDE (103) partner repository with the dataset identifiers: PXD052437 for Total protein extracts and PXD052348 for IP eluate fraction. https://www.ebi.ac.uk/pride/login.

## Acknowledgements

The authors would like to thank the following people for their help and support: Eduardo Muñoz-Díaz for advice on FISH experiments, Claire Picart for help with confocal microscopy, Arnaud Dannfald for help and advice with polysome profiling, Valentin Carriere for help with IP experiments, Sandrine Koechler from the IBMP gene expression analysis platform, Valérie Cognat from the IBMP bioinformatics platform, Christina Berrissou for Mim-tRNA-seq experiments and Julien Henry from the GenoToul bioinformatics facility. We would also like to thank Guillaume Née for his insightful and productive discussions on seed germination and seedling establishment.

## Author contributions

Molecular, biochemical and genetics experiments: SAN and JAF. Immunolocalisation experiments: PC; RT-PCR and technical assistance: AdB; 5’Pseq analysis: MCC and RM; LC MS/MS analyses: MR and CC; bioinformatics analysis: JH, MCC, and CG; RiboMethSeq analyses: VM, VM and YM; Mim-tRNA-seq analysis: LD; supervising the project: JSV, YM and CG; analysing the data: JSV, YM, CG and SAN; wrote the manuscript: SAN and JSV. All authors approved the final manuscript.

